# Riboflavin Depletion Promotes Longevity and Metabolic Hormesis in *Caenorhabditis elegans*

**DOI:** 10.1101/2022.06.30.498343

**Authors:** Armen Yerevanian, Luke Murphy, Sinclair Emans, Yifei Zhou, Fasih Ahsan, Daniel Baker, Sainan Li, Adebanjo Adedoja, Lucydalila Cedillo, Einstein Gnanatheepam, Khoi Dao, Mohit Jain, Irene Georgakoudi, Alexander Soukas

## Abstract

Riboflavin is an essential cofactor in many enzymatic processes and in the production of flavin adenine dinucleotide (FAD). Here we report that the partial depletion of riboflavin through knockdown of the *C. elegans* riboflavin transporter 1 (*rft-1*) promotes metabolic health by reducing intracellular flavin concentrations. Knockdown of *rft-1* significantly increases lifespan in a manner dependent on FOXO/*daf-16*, AMP-activated protein kinase (AMPK)/*aak-2*, the mitochondrial unfolded protein response, and mTOR complex 2 (mTORC2). Riboflavin depletion promotes altered energetic and redox states and increases adiposity, independent of lifespan genetic dependencies. Riboflavin depleted animals also exhibit activation of caloric restriction reporters without a reduction in TORC1 signaling. Our findings indicate that riboflavin depletion activates an integrated, hormetic response that promotes lifespan and healthspan in C. elegans.

## Introduction

Healthy mitochondrial function requires the coordination of multiple cellular inputs including sufficient energetic substrates, amino acids and micronutrients. Vitamin cofactors are key to metabolic processes such as the citric acid cycle, electron transport chain physiology, and energy shuttling into the cytosol. One family of vitamins that participate in mitochondrial physiology are the flavins (Mansoorabadi, Thibodeaux et al. 2007). The flavin co-factors include flavin mononucleotide (FMN) and flavin adenine dinucleotide (FAD) and are essential for redox chemistry and electron shuttling (Massey 1995). FAD is classically known to serve as an electron acceptor in the conversion of succinate to fumarate by succinate dehydrogenase in the citric acid cycle, as well as an electron donor to complex II of the electron transport chain. The flavins are also cofactors for multiple enzyme classes including the oxidoreductases and the fatty acid dehydrogenases (Lienhart, Gudipati et al. 2013). They are derived from riboflavin, a water soluble ribitol derivative also known as vitamin B2. Riboflavin is an essential nutrient for all animals that must be acquired either from food sources or from commensal gut flora (Powers 2003).

The animal kingdom has evolved specific transporters to import riboflavin from the gut lumen and to transport them intracellularly. (Moriyama 2011) In humans and mice, three isoforms of these transporters are expressed by three distinct genes: Slc52A1, Slc52A2 and Slc52A3 (Yonezawa, Masuda et al. 2008, Yamamoto, Inoue et al. 2009, Subramanian, Subramanya et al. 2011). Disruption in the function of these transporters is known to induce pathology through cellular riboflavin depletion (Nabokina, Subramanian et al. 2012). Congenital deficiency in these transporters is associated with clinical syndromes in humans including Brown-Vialetto-Van Laere syndrome, where patients experience progressive neurologic deficits, and is treated with extremely large doses of riboflavin (Dipti, Childs et al. 2005, Spagnoli and De Sousa 2012). Nutritional riboflavin deficiency (ariboflavinosis) is a very rare but also described clinical disorder which is associated with dermatologic and hematologic manifestations (Sydenstricker 1941).

Despite the clinical and epidemiologic data describing syndromes of riboflavin deficiency, the metabolic impacts of riboflavin depletion at a cellular level are not well described. Flavin deficiency is thought to alter fatty acid metabolism through disrupted beta-oxidation as well as reduced flux through the citric acid cycle due to a lack of FAD. We hypothesized that these disruptions may lead to the activation of signaling pathways that are associated with mitochondrial and energetic stress. We further hypothesized that partial depletion of riboflavin will have beneficial metabolic effects by activating cellular responses associated with caloric restriction and longevity.

In the present study, we determine that physiologic riboflavin depletion alters cellular energetics and activates key longevity factors such as AMPK, FOXO, and the mitochondrial unfolded protein response (UPR^mt^). We report that riboflavin depletion via riboflavin transporter knockdown extends lifespan and promotes healthspan in *C. elegans*, suggesting that riboflavin depletion provides metabolic benefits and mimics favorable features of caloric restriction without actual reductions in caloric intake.

## Results

### Knockdown of *rft-1* Promotes Longevity via Riboflavin Depletion

We elected to approach the hypothesis that partial riboflavin depletion may provide metabolic benefits by performing functional genomics in *C. elegans*. The nematode genome encodes two riboflavin transporter orthologs, *rft-1* and *rft-2* (Biswas, Elmatari et al. 2013). *rft-1* exhibits strong intestinal expression, so we first examined consequences of its knockdown (Gandhimathi, Karthi et al. 2015). *rft-1* depletion via RNA interference (RNAi) prompts a significant, 25% increase in lifespan (**Figure 1a**). The RNAi knockdown of *rft-1* reduces the transporter’s mRNA by approximately 70%, suggesting that partial riboflavin transporter deficiency rather than complete knockdown induces this phenotype, as *rft-1* mutants are not viable (**Figure 1b**). mRNA encoding the paralogous riboflavin transporter *rft-2* trends non-significantly upwards with *rft-1* RNAi, which indicates that there is not significant compensation for *rft-1* knockdown by *rft-2*, and confirms the specificity of the RNAi-based knockdown of *rft-1* (**Figure 1b**). To confirm that the longevity phenotype of the *rft-1* knockdown animals is due to reduced riboflavin uptake rather than a non-canonical effect of the transporter, we administered high dose riboflavin supplementation to attempt to overcome the deficit in transport. As expected, high doses of riboflavin abrogate the lifespan increase attributable to *rft-1* RNAi (**Figure 1a**). This strongly suggests that riboflavin is the etiologic factor in the *rft-1* RNAi phenotype, and further that depletion of riboflavin due to transporter deficiency is the source of lifespan extension. Worm lysates of young adult worms treated with *rft-1* RNAi exhibit marked reductions in riboflavin, FMN and FAD levels as assessed by quantitative liquid chromatography/mass spectrometry (LC/MS) **(Figure 1c)**. In parallel, we evaluated the expression of enzymes key to flavin co-enzyme synthesis including riboflavin kinase (*rfk-1*) and FAD synthetase (*flad-1*). *rft-1* RNAi was not associated with reductions in expression of *flad-1* and *rfk-1* which produce FAD and FMN respectively, suggesting stoichiometric depletion of FMN/FAD rather than reductions in the enzymes that govern production **(Figure S1a)**.

**FIGURE 1.**
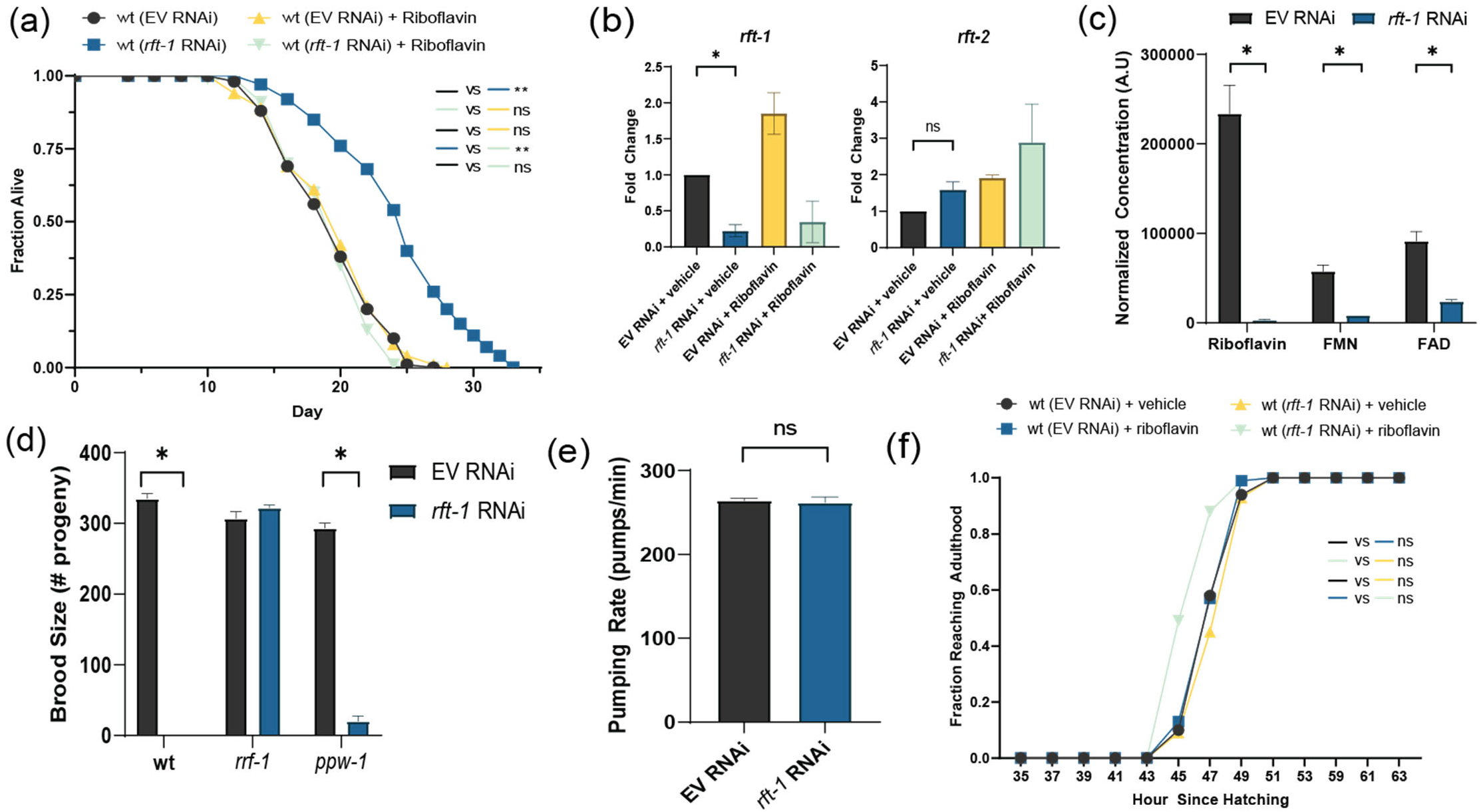
*rft-1* RNAi depletes flavins and extends lifespan. (a) Knockdown of the riboflavin transporter *rft-1* via RNA interference significantly extends lifespan in wild type (wt) animals versus empty vector RNAi (EV). Addition of 665 μM riboflavin abrogates this lifespan extension (see table S1 for tabular data and replicates). (b) Knockdown of *rft-1* produces 70% reduction in transcript levels by quantitative RT-PCR and does not induce corresponding reductions in orthologous transporter *rft-2* mRNA. (c) LC-MS/MS analysis of worm lysates collected at adult day 1 treated with *rft-1* RNAi reveals significant reductions in organismal riboflavin, FMN, and FAD concentrations. (d) Brood size is diminished in *rft-1* RNAi treated animals and remains suppressed in *ppw-1* animals (somatic RNAi competent, germline RNAi incompetent) but is rescued in *rrf-1* animals (somatic RNAi blunted, germline RNAi competent), suggesting a somatic site of *rft-1* action. (e) Pharyngeal pumping assay on adult day 1 animals reveals no difference between EV and *rft-1* RNAi. (f) Developmental rate from the first larval stage to adulthood is unchanged in animals treated with *rft-1* RNAi, with or without riboflavin. For a-f, results are representative of three biological replicates. * indicates *P* < 0.05, **, *P* < 0.001 by log-rank analysis (a and f), two-way ANOVA of ΔCt values (b), standard two-way ANOVA (c-d) and two-tailed Student’s t-test (e). Bars represent means ± SEM. Ns presents non-significant.

Previous descriptions of brood size deficits in *rft-1* knockdown animals suggests that the germline may play a role in the phenotype(Biswas, Elmatari et al. 2013, Qi, Kniazeva et al. 2017, Yen, Ruter et al. 2020). We utilized *rrf-1* (somatic RNAi blunted, germline RNAi competent) and *ppw-*1 (somatic RNAi competent, germline RNAi incompetent) mutants and examined brood size on *rft-1* RNAi. The loss of brood size is dependent on somatic action of *rft-1* (i.e. normalized in *rrf-1* mutants which do not efficiently conduct somatic RNAi). This suggests that a global process is altering metabolic and reproductive capacity **(Figure 1d)**. The reduction in brood size raised the question of whether germ line stem cells and oocyte production is altered by riboflavin depletion. We examined the presence of the germline stem cells and oocytes via DAPI staining, which revealed an intact germline and oocyte production (**Figure S1b**). Riboflavin deficiency is associated with neurologic sequelae in mammals, and we wanted to verify that feeding behavior is unchanged compared to control animals(Qi, Kniazeva et al. 2017). Knockdown of *rft-1* does not affect eating behavior as measured by the pharyngeal pumping rate (**Figure 1e**). We also sought to evaluate whether there was a change in developmental rate as a potential source for this phenotype. RNAi to *rft-1* does not significantly change the rate of development to adulthood. RNAi of *rft-1* in the RNA interference sensitive strain *eri-1* also does not alter developmental rate **(Figure 1f)**. These findings suggest that animals treated with *rft-1* knockdown exhibit normal development and behavior, and that nutritional depletion of riboflavin does not preclude them from reaching adulthood.

### Riboflavin Depletion Promotes Longevity Through FOXO

FOXO is known to act genetically downstream of nutrient-sending manipulations that extend lifespan (Lee, Kennedy et al. 2003, Greer, Dowlatshahi et al. 2007) and we hypothesized that it may be activated in the context of riboflavin depletion. Indeed, *daf-16* loss of function is epistatic to lifespan extension with *rft-1* RNAi, with or without the presence of supplemental riboflavin **(Figure 2a)**. A DAF-16::GFP translational reporter demonstrates greater nuclear localization at adult day 1 and day 3 with riboflavin depletion, suggesting that *rft-1* RNAi activates DAF-16 **(Figure 2b and S2a)**. Transgenic DAF-16::GFP worms treated with empty vector and *rft-1* RNAi as synchronous L1 were subsequently transferred at adult day 1 to plates with and without riboflavin. By Adult Day 3, additional riboflavin completely abrogates the DAF-16 nuclear localization evident in in *rft-1* RNAi treated animals **(Figure S2b)**. This indicates that the consequences of riboflavin depletion on *daf-16* nuclear localization are reversible in adulthood. Confirming increased transcriptional activity of DAF-16 in the setting of riboflavin depletion, RNAi of *rft-1* led to the upregulation of a *sod-3p*::GFP reporter **(Figure 2c)**. This activation was present through adult day 7, indicating consistent FOXO activation post-developmentally **(Figure S2c)**.

**FIGURE 2.**
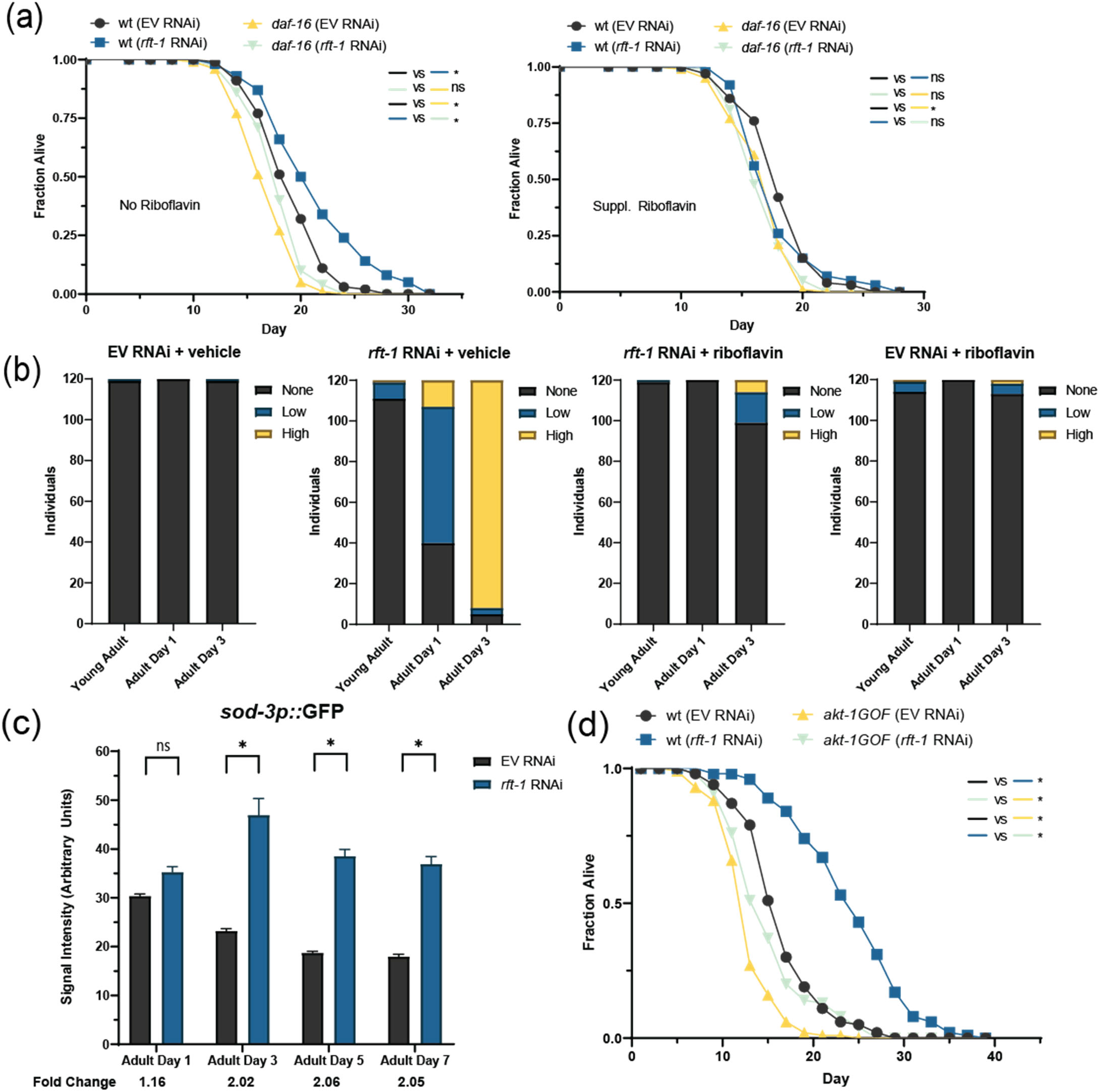
Riboflavin depletion promotes longevity by activating FOXO/*daf-16*. (a) Lifespan extension with knockdown of the *rft-1* transporter is abrogated in *daf-16* mutants, with and without supplemental riboflavin. (b) Nuclear localization of DAF-16 was increased following *rft-1* RNAi in a riboflavin-dependent manner as assessed by a DAF-16::GFP translational reporter. Animals were binned into no nuclear localization, low levels of localization and high levels of localization (see Fig. S2 for representative images). N = 60 animals per condition, representative of two biological replicates. (c) A *sod-3p*::GFP transcriptional reporter indicates increased activity of DAF-16 significantly over the lifespan of *rft-1* RNAi treated animals. (d) A chromosomally-located *akt-1* gain-of-function mutation blunts the longevity response to riboflavin depletion vs. wt. For a-d, results are representative of at least two biological replicates. * indicates *P* < 0.05 by log-rank analysis (a and d), and by two-way ANOVA followed by Sidak’s multiple comparisons post-hoc test (c). See table S1 for tabular data and replicates of survival analyses. Bars represent means ± SEM.

The activation of FOXO suggests either suppression of insulin-like/PI-3 kinase signaling (canonical FOXO activation) or activation via another mechanism. We examined whether insulin signaling was playing a role by examining the effect of *rft-1* RNAi on *akt-1* and *pdk-1* gain of function mutants (Paradis and Ruvkun 1998). These animals are short lived due to constitutive inhibition of FOXO. The *akt-1* gain-of-function mutation abrogates lifespan extension attributable to *rft-1* RNAi **(Figure 2d)**. The *pdk-1* gain-of-function mutant on *rft-1* RNAi, conversely, still exhibits lifespan extension **(Figure S2d)**. This indicates that suppression of DAF-16 via Akt abrogates lifespan extension prompted by riboflavin deficiency.

### Riboflavin Depletion Alters Cellular Redox Ratio and Energetics

We suspected that AMPK may be similarly activated by energetic stress with riboflavin deficiency, and further that AMPK activation may be mechanistically linked to lifespan extension with *rft-1* RNAi. Knockdown of *rft-1* fails to promote longevity with loss of function in the AMPKα catalytic subunit *aak-2* (**Figure 3a**), and this pattern is not altered by addition of riboflavin **(Figure 3b**). Consistent with AMPK activation by riboflavin depletion, phospho-AMPK^T172^ levels are increased in *rft-1* RNAi treated animals, and this effect is abrogated by the addition of riboflavin **(Figure 3c)**. In order to determine whether AMPK is required for activation of DAF-16 under riboflavin depletion, we examined DAF-16::GFP nuclear localization with and without functional *aak-2*. Under *rft-1* RNAi conditions, DAF-16 nuclear localization still increases in the absence of *aak-2* **(Figure S3a)**. This could suggest that AMPK and FOXO are each required in parallel to promote lifespan downstream of riboflavin deficiency.

**FIGURE 3.**
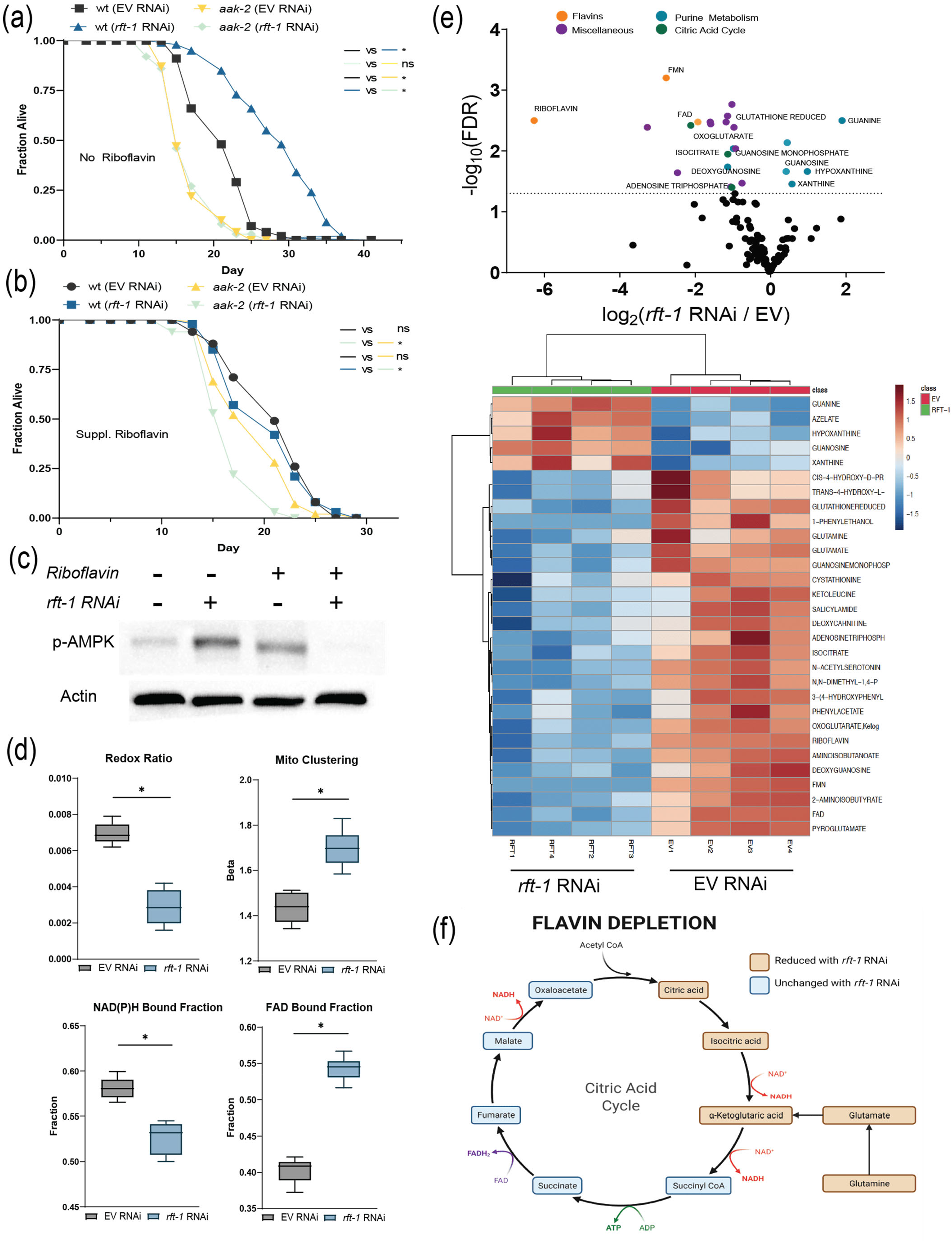
Riboflavin depletion alters cellular energetics. (a) Lifespan extension with *rft-1* RNAi is abrogated in AMPK/*aak-2* mutants. (b) Addition of riboflavin has a deleterious effect on lifespan in *aak-2* mutants. (c) Western blotting of phospho-AMPK^Thr172^ in lysates collected from young adult animals indicates activation following RNAi to *rft-1*, an effect abrogated by the addition of riboflavin. (d) Box plots of results from two-photon and fluorescence lifetime imaging, including organismal redox ratio, mitochondrial clustering, NAD(P)H bound fraction and FAD bound fraction for EV and *rft-1* RNAi treated animals. Riboflavin depletion decreases the redox ratio and increases intestinal mitochondrial clustering and the FAD bound fraction. (e) Volcano plot and heatmap of differentially abundant metabolites quantified by LC-MS reveals reductions in citric acid metabolites including citric acid, isocitric acid and α-ketoglutarate following riboflavin depletion. Purine metabolites including xanthine, hypoxanthine and guanosine are enriched with *rft-1* RNAi. (f) Representation of citric acid metabolites impacted by riboflavin depletion. See table S1 for tabular data and replicates of survival analyses. For (a-c) results are representative of two biological replicates. For (d) results are from 8 worms of two biological replicates. * indicates *P* < 0.01 by log-rank analysis (a-b) and by two tailed t-test (d). Box and whisker plots (d) indicate median and 5/95^th^ percentile respectively.

Activation of AMPK suggests that modulation of cellular energetics might play a role in the longevity phenotype seen with the *rft-1* knockdown. We hypothesized that reductions in flavin cofactor (FAD, FMN) concentrations induce mitochondrial stress responses due to changes in organellar energetics by altering redox state. We examined the impact of riboflavin depletion on the redox ratio utilizing label-free multiphoton microscopy and fluorescence lifetime imaging (FLIM) of intestinal cells in control and *rft-1* RNAi treated animals **(Figure Sb-c)**. Animals treated with *rft-1* RNAi versus empty vector control-treated animals exhibit a significant decrease in the optical redox ratio, defined as the ratio of FAD/(NAD(P)H + FAD) calculated based on the autofluorescence signatures of the corresponding co-enzymes. There is also an increase in mitochondrial clustering, suggesting altered mitochondrial energetics in an oxidized state and morphologic changes to the mitochondrial network **(Figure 3d)**(Liu, Pouli et al. 2018). A significant decrease in the NAD(P)H protein bound fraction suggests decreased levels of glutaminolysis and enhanced utilization of the glutathione pathway (Alonzo, Karaliota et al. 2016). An increase in the FAD bound fraction suggested that overall FAD depletion was causing aggressive capture of flavin co-factors by enzymatic machinery **(Figure 3d)**. LC/MS metabolomics of *rft-1* RNAi treated animals indicates significant changes in multiple metabolic pathways. Increases in purine catabolism metabolites were present, including xanthine, hypoxanthine, and guanosine **(Figure 3e)**. Pathway enrichment analysis reveals that other than riboflavin metabolism, riboflavin deficiency leads to significant impact on metabolite changes in the glutathione and purine metabolism pathways **(Figure S3d)**. Components of the citric acid cycle including citrate, isocitrate, and α-ketoglutarate, as well as ATP, were reduced by riboflavin depletion. Glutamate and glutamine levels were also reduced suggestive of disruptions in glutamine synthesis **(Figure 3f)**.

### Riboflavin Depletion Activates the Mitochondrial Unfolded Protein Response

The frank changes in energetic status, altered redox ratio, and presence of mitochondrial clustering all suggested that mitochondrial stress responses may also be contributing to the longevity response to riboflavin deficiency. Indeed, the UPR^mt^ is activated by *rft-1* knockdown, as evidenced by induction of *hsp-6p*::GFP (Labbadia, Brielmann et al. 2017) induction on days 1 and 3 of adulthood, and this effect is abrogated by the addition of riboflavin **(Figure 4a)**. Full activation of the UPR^mt^ is known to require the transcription factor ATFS-1, which translocates to the nucleus to activate stress response pathways (Wu, Senchuk et al. 2018). Established target genes of ATFS-1, including *cdr-2, hrg-9*, and *C07G1*.*7* (Soo and Van Raamsdonk 2021) are upregulated with *rft-1* knockdown, with the previously undescribed gene C07G1.7 exhibiting a 2000-fold increase **(Figure 4b)**. The UPR^mt^ activation is necessary for lifespan extension, as a*tfs-1* loss of function animals, which have lower lifespans compared to wild-type at baseline, do not exhibit lifespan extension with *rft-1* knockdown **(Figure 4c)**. In the setting of gain-of-function mutations in *atfs-1*, which lead to shortened lifespan (Bennett, Vander Wende et al. 2014), *rft-1* knockdown still promotes significant extension of lifespan **(Figure 4d)**. This indicates that the UPR^mt^ response is necessary but not sufficient for the riboflavin depletion longevity phenotype.

**FIGURE 4.**
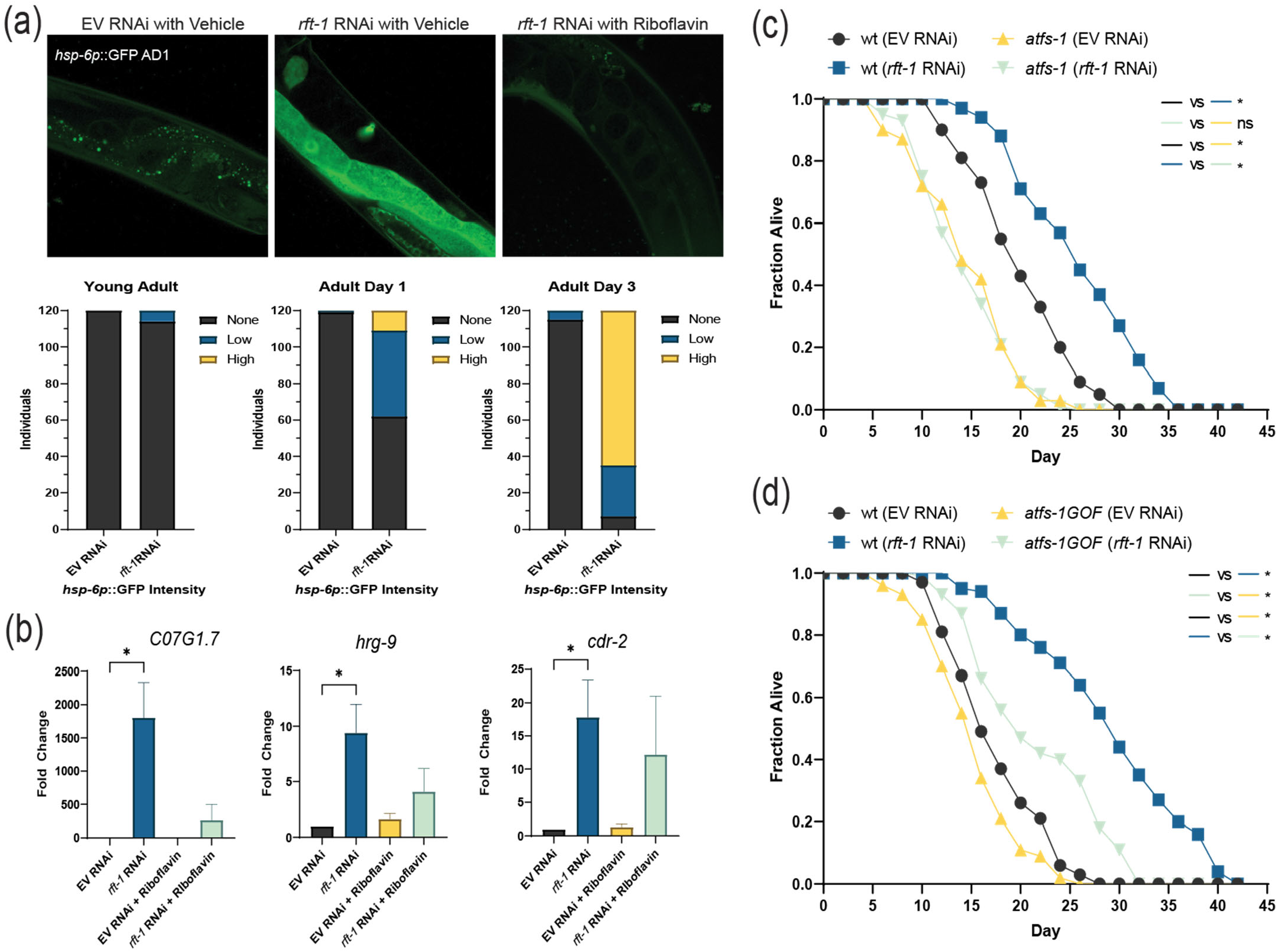
Activation of the mitochondrial unfolded protein response (UPR^mt^) is required for riboflavin depletion to promote longevity. (a) RNAi to *rft-1* promotes activation of *hsp-6*p::GFP progressively on adult days 1 and 3, an effect reversed by riboflavin supplementation (for binning, N = 60 worms per condition, representative of two biological replicates) Images above at 40X, binning performed at 10X magnification. (b) Quantitative RT-PCR of established *atfs-1* target genes indicates marked upregulation with riboflavin depletion. (c) *atfs-1* loss of function mutants do not experience lifespan extension with riboflavin depletion. (d) Gain of function mutants in *atfs-1* are short lived but preserve responsiveness to *rft-1* RNAi. For (b), results are representative of at least three biological replicates unless otherwise specified. See table S1 for tabular data and replicates of survival analyses. * indicates *P* < 0.05 by two-way ANOVA of ΔCt values (b), and by log-rank analysis (c and d). Bars represent means ± SEM.

### Riboflavin Depletion Alters Somatic Lipid Stores

The long-lived phenotype of riboflavin depletion and the role of flavin cofactors in beta-oxidation suggests that changes in lipid composition may be manifest following *rft-1* knockdown. We hypothesized that changes in lipid metabolism occur upstream of or in parallel to FOXO activation, due to changes in enzymatic function (such as reduced activity of lipid dehydrogenases). RNAi of *rft-1* induces significant increases in fat mass in both the intestine and the germline in adult day 1 worms, as exhibited by fixative oil-red-O and Nile red staining **(Figure 5a, S5a)** Epistasis analysis indicates that *rft-1* RNAi increases fat mass in *daf-16, aak-2* and *raga-1* animals but not in *atfs-1* or *rict-1* animals by fixative-based Nile red staining **(Figure 5a)**. Post-developmental *rft-1* RNAi beginning at young adult stage in enhanced RNAi *eri-1* animals also increases fat mass, indicating that riboflavin deficiency does not impact lipid metabolism exclusively through a developmental pleiotropy **(Figure 5a)**. Confirming these observations and further delineating the nature of the lipids increased in abundance following *rft-1* RNAi, stimulated Raman scattering (SRS) analysis of live adult day 1 animals indicates increased total signal of unsaturated fatty acids and the unsaturated to total lipid ratio in riboflavin deficiency **(Figure S5b)** (Nieva, Marro et al. 2012, Potcoava, Futia et al. 2014). By gas chromatography/mass spectroscopy (GC/MS) of triglyceride and phospholipid fractions separated by solid phase extraction, global triglyceride stores increase by 40% in both young adult and adult day 1 *rft-1* RNAi-treated animals, consistent with the spectroscopic imaging and fixative-based lipid staining **(Figure 5b)**. While only small changes are evident by young adulthood **(Figure S5c)**, by adult day 1 animals exhibit significant differences in their lipid composition, with increases in unsaturated and branched chain fatty acids, and reductions in cyclopropyl fatty acids in both triglyceride and phospholipid fractions **(Figure 5c, S5c)**. Due to increases in branched chain fatty acid synthesis, we examined the expression of *acdh-1*, which is a known branched chain dehydrogenase in *elegans* and that has been previously reported as a dietary sensor (Watson, MacNeil et al. 2013). An *acdh-1* promoter GFP reporter is significantly increased ∼70% with *rft-1* RNAi at adult day 1 **(Figure 5d)**. In order to begin to determine whether unsaturated fatty acids are elevated in riboflavin deficiency owing to increased production vs utilization, we examined expression of the fatty acid desaturase *fat-7* (Han, Schroeder et al. 2017). Riboflavin depletion does not promote changes in *fat-7* expression early in life but preserves it with aging **(Figure S5d)**.

**FIGURE 5.**
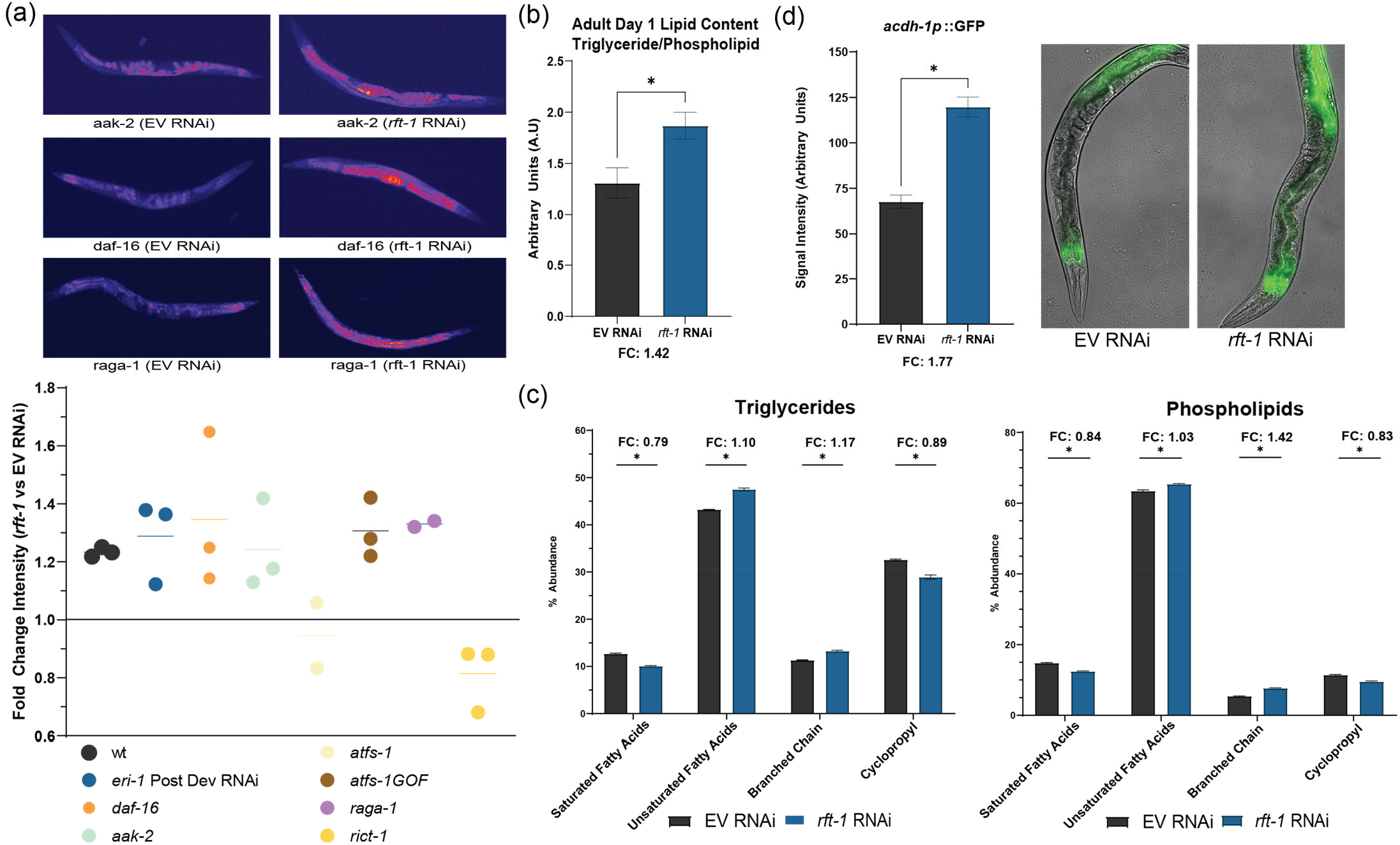
Riboflavin depletion alters lipid metabolism. (a) Fixative-based Nile red staining and quantification of images reveal significant increases in fat mass with post developmental *rft-1* RNAi in *eri-1* enhanced RNAi mutants and with larval exposure to *rft-1* RNAi in multiple mutants including *daf-16, aak-2, atfs-1(GOF)*, and *raga-1. atfs-1* and *rict-1* loss-of-function mutants do not exhibit increased fat on *rft-1* RNAi. (b) Lipid analysis via GC/MS reveals an increase in overall fat stores (triglyceride/phospholipid ratio). (c) Shifts towards unsaturated fatty acid side chains and branched chain lipids in phospholipid and triglyceride fractions in aggregate (d) *acdh-1p*::GFP expression is significantly upregulated in intestine with *rft-1* RNAi. * indicates *P* < 0.05 by students 2-tailed t-test (b,d) and by two-way ANOVA (c). Bars (b-d) represent means ± SEM. Dots in (a) represent individual biological replicates.

### Riboflavin Depletion Activates Dietary Restriction Pathways

The long-lived phenotype of riboflavin depletion, concomitant with decreases in energetics, AMPK activation, and impairment in lipid beta-oxidation, suggested to us that riboflavin depletion resembles a dietary restriction-like phenotype. The *acs-2* and *bigr-1* genes have been well established to be transcriptionally upregulated during periods of caloric restriction in *C. elegans (Van Gilst, Hadjivassiliou et al. 2005, Wu, Zhou et al. 2016). rft-1* RNAi induced *bigr-1*::RFP and *acs-2p*::GFP expression with age **(Figure 6a)**. We sought to assess whether other canonical caloric restriction factors were involved in riboflavin depletion, particularly the *C. elegans* FoxA homolog *pha-4* (Panowski, Wolff et al. 2007). Consistent with our hypothesis, lifespan extension with *rft-1* RNAi is dependent on *pha-4*/*FoxA* **(Figure 6b)**. Inhibition of target of rapamycin (TOR) signaling is also important in the response to dietary restriction. Thus, we examined whether mutants in the TOR complex 1 (TORC1) and TOR complex 2 (TORC2) pathways exhibit longevity with riboflavin depletion. RAGA/*raga-1* mutants experience lifespan extension on *rft-1* RNAi **(Figure 6c)**. To further determine whether altered TORC1 activity is required for the hormetic effects of riboflavin depletion, we used a strain of elegans that contains a knock-in, humanized S6K, which permits immunoblotting for phospho-S6K to determine the activity of TORC1. No difference is evident in p-S6K between *rft-1* RNAi and empty vector control, suggesting that altered TORC1 signaling is not essential to the biological response to riboflavin depletion **(Figure 6d)**. In contrast, loss of function mutations in the essential TORC2 subunit *rict-1* experience no lifespan extension with *rft-1* RNAi, indicating that riboflavin depletion requires TORC2 activity to exact its favorable effects **(Figure 6e)**.

**FIGURE 6.**
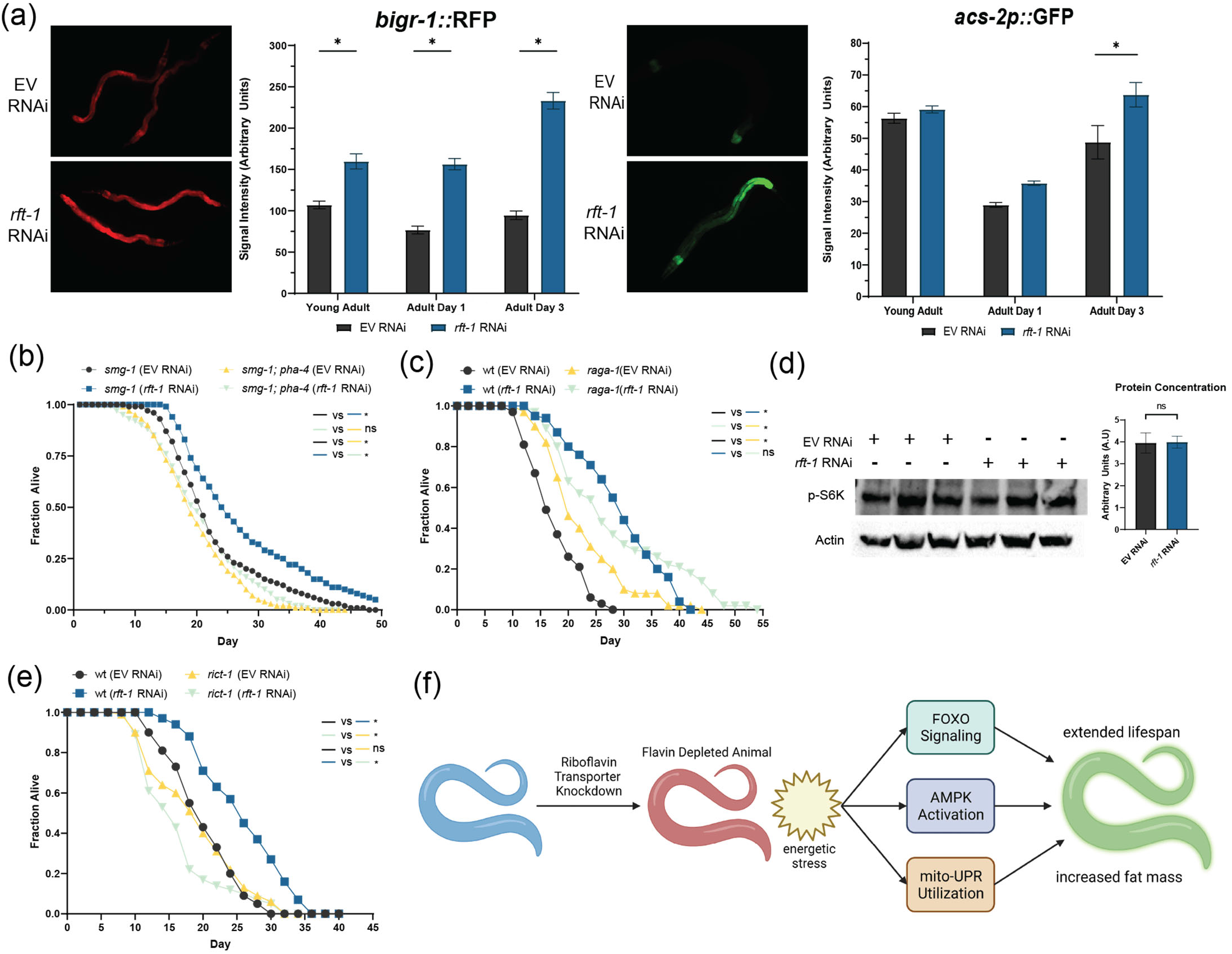
Riboflavin depletion mimics features of dietary restriction. (a) Imaging of reporters known to be activated with dietary restriction (*bigr-1*::RFP, *right*, and *acs-2p*::GFP, *left*) indicates activation under *rft-1* RNAi that progresses with age. (b) Lifespan extension with *rft-1* RNAi is abrogated in *pha-4* loss-of function mutants versus temperature sensitive *smg-1* mutant controls. (c) TORC1 mutant *raga-1* exhibits lifespan extension with *rft-1* RNAi. (d) Western blot of adult day 1 animals containing humanized S6K (permitting western blotting for TORC1-mediated phosphorylation of S6K^T389^) reveals no change in phospho-S6K concentrations between control and riboflavin depleted worms. (e) TORC2 mutant *rict-1* does not exhibit lifespan extension with riboflavin depletion. (f) Model of riboflavin depletion impact on metabolism and longevity. Knockdown of the riboflavin transporter leads to depletion of flavin co-factors, influencing the energetic status of the animal and affecting global enzymatic activities dependent on FAD and FMN. This creates an integrated metabolic signaling response that promotes increased triglyceride stores and extends lifespan. See table S1 for tabular data and replicates of survival analyses. * indicates *P* < 0.05 by two-way ANOVA (a), log-rank test (b,c and e). Bars represent means ± SEM

## Discussion

Vitamins, as essential co-factors for life, have traditionally been viewed as highly beneficial entities independent of their concentrations. This is particularly true of the B-class vitamins which are water soluble and do not exhibit significant toxicities at moderate supraphysiologic doses. Our work in *C. elegans* counters the notion that more is always better, as depletion of key enzymatic co-factor riboflavin can trigger metabolic and physiologic stress responses that are hormetic in nature and extend lifespan.

We were initially concerned that the reduction of the flavin cofactors such as FAD and FMN would be frankly toxic to the organism. This was particularly true when LC/MS revealed that FAD levels in the *rft-1* treated animals were 80-90% below normal. Alternatively, it is clear from data presented here that this level of reduction promotes energetic perturbations most consistent with disruptions in mitochondrial respiration. Unexpectedly, however, the riboflavin depletion phenotype is different than that of classic electron transport chain (ETC) disruption, such as in *cco-1* and *frh-1* (frataxin) knockdowns (Durieux, Wolff et al. 2011, Schiavi, Torgovnick et al. 2013). Previous examinations of *cco-1* and *frh-1* RNAi have shown that lifespan extension with ETC disruption is AMPK and FOXO independent (Durieux, Wolff et al. 2011, Schiavi, Torgovnick et al. 2013). Unlike traditional ETC disruption, riboflavin depletion’s lifespan extending properties depend on these key energetic sensing pathways. We anticipated that intestinal FOXO activation would depend on increased AMPK activity, however nuclear localization of DAF-16 still occurs in AMPK mutants, indicating AMPK-independent governance of FOXO activity. Alternatively, FOXO activation may be dependent upon inhibition of *akt* and/or insulin-like signaling as gain-of-function mutations in *akt*-*1* also suppress lifespan prolongation by riboflavin deficiency. Why this phenotype is not shared with *pdk-1* gain-of-function mutations could be because of the relative strength of the latter, or complexities of riboflavin deficiency on cellular signaling. Determining between these possibilities will requires additional testing. Finally, additional requirements of the UPR^mt^ and potentially TORC2 for lifespan extension suggest that these pathways function as parallel, critical effectors to achieve a concerted cellular response to the reductions of FAD and FMN **(Figure 6f)**.

Riboflavin depleted animals exhibit normal food consumption (pharyngeal pumping rate), normal TORC1 activation, and elevated triglyceride stores, suggesting that the phenotype is not due to a reduction in macronutrient availability, but may be due to defective nutrient mobilization and utilization. The dependencies on FOXO/*daf-16*, FOXA/*pha-4*, and AMPK/*aak-2*, as well as activation of canonical “starvation” reporters *acs-2* and *bigr-1*, suggest that riboflavin depleted animals experience a caloric restriction-like phenotype with micronutrient depletion only. Amino acid sensing and dietary restriction via essential nutrients such as methionine have been previously described to extend lifespan (Cabreiro, Au et al. 2013). Depletion of canonical vitamin co-factors however, has proven deleterious in previous investigations. Depletion of biotin, B12 derivatives, and folate have previously shown to shorten lifespan (Austin, Liau et al. 2010, Bito, Matsunaga et al. 2013). To our knowledge, our work is the first to show that depletion of a vitamin co-factor can mimic features of dietary restriction and extend lifespan using shared molecular mechanisms.

Elevated triglyceride stores following riboflavin depletion is independent of canonical lifespan regulating pathways such as FOXO, AMPK and TORC1. This decoupling of fat mass and lifespan suggest that the lipid phenotype may be regulated by enzymatic processing of lipids upstream of the energetic stress axes. The exceptions to this were the *atfs-1* and *rict-1* mutants. Phenotypes associated with UPR^mt^ activation are known to induce lipid accumulation (Kim, Grant et al. 2016, Yang, Li et al. 2022). Recent work has identified NHR-80 as a key regulator of citrate sensing and lipid accumulation in the UPR^mt^ phenotype (Yang, Li et al. 2022). The relevance of *atfs-1* activity to *dve-1* and *ubl-5* function suggests that the UPR^mt^ may be instrumental in the communication of flavin depletion and related citric acid cycle disruptions on organismal energetics. The lack of fat mass increase in *rict-1* mutants, which at baseline exhibit higher lipid content, suggests either a dependency or inability for riboflavin depletion to overcome the excess lipid accumulation associated with TORC2 knockout. TORC2 has been well described as a nutritional sensor that regulates lipid biogenesis (Soukas, Kane et al. 2009), and it is entirely plausible that there are distinct inputs in mitochondrial energetics, mito-stress and TORC2 activation that are governed by flavin biology. Further investigation would be beneficial to identify whether TORC2 can directly sense changes in flavin levels, as this would have significant implications for the nutritional regulation of anabolic signals in senescence and cancer.

*rft-1* RNAi does not impact developmental rate and metabolic phenotypes manifest most impressively at the young adult to adult day 1 transition. This is in parallel to the growth and development of the germline and the oocytes. This pattern suggests either that the larval stages are relatively resistant to riboflavin depletion, or, more likely, contain and accumulate sufficient flavin cofactors at time of egg lay and during early larval development (prior to *rft-1* knockdown) to proceed through development normally. We surmise that somatic growth dilutes endogenous flavin cofactors, subsequently inducing the favorable effects of riboflavin deficiency uniquely in adulthood. Further, development of the germline and riboflavin shunting into oocytes in late larval and early adult stages may prompt further, rapid riboflavin depletion, inducing the metabolic stress required to induce the phenotype identified in this work. It is worthy of mention that riboflavin deficiency leads to sterility, but this sterility is not accompanied by a decrease in germline stem cell numbers. Thus, effects on the germline are unlikely to be responsible for the shifts in lifespan and fat mass evident in riboflavin deficiency.

The presence of a post-developmental fat increase with depletion of riboflavin suggests that acute depletion in adulthood has important impacts that are likely different from depletion starting at larval stages. The most likely etiology for these post-developmental changes are alterations in enzymatic activity due to loss of flavin co-factors. The flavin co-factors are important for a wide variety of enzymes particularly those associated with oxidoreductase functions including the fatty acid dehydrogenases. The ‘flavoproteome’ is an established set of enzymes requiring FAD, FMN or riboflavin to function (Lienhart, Gudipati et al. 2013). The impact of riboflavin depletion globally on the proteome is likely to produce stoichiometric shifts in key metabolites that will alter global physiology. Differential utilization of different dehydrogenases (branched chain vs long chain) may also explain the unique lipid phenotype that is produced with riboflavin depletion. Using metabolomics, we identified other examples of likely enzymatic effects, with evidence of reductions in purine catabolism likely due to loss-of-function in xanthine dehydrogenase. Alterations in xanthine metabolism have been previously described as beneficial and lifespan extending (Gioran, Piazzesi et al. 2019). The impact of riboflavin depletion on enzymatic processes is complex and there are likely to be both hormetic and harmful impacts of this process. Identifying the enzymatic pathways where riboflavin depletion provides beneficial versus detrimental impacts will provide new insights into mechanistic targets promoting longevity. We suggest that further investigations into the functions of the flavoproteome and flavin biology will serve to identify new therapeutic and investigational targets for the metabolism of aging and aging associated diseases.

## Experimental Methods

### *C. elegans* genetics

Strains were maintained at 20°C on nematode growth medium (NGM) plates seeded with E. coli OP50. All experiments were conducted at 20°C unless otherwise specified. The following strains were utilized: wild type (N2 Bristol ancestral), NL3511 *ppw-1*(*pk1425*), NL2098 *rrf-1*(*pk1417*), *daf-16*(*mgDF47*), TJ356 zls356[daf-16p::daf-16a/bGFP+rol-6(su1006), CF1553 *muls84*[(pAD76)*sod-3p*::GFP+*rol-6*(*su1006*)], GR1318 *pdk-1*(*mg142gf)*, GR1310 *akt-1*(*mg144gf*), RB754 *aak-2*(*ok524*), VC3201 *atfs-1*(*gk3094*), QC118 *atfs-1*(*et18*), SJ4100 *zcls13*[*hsp-6p*::GFP+*lin-15*(+)], DMS303 *nls590*[fat-7p::fat-7::GFP +lin15(+)], VL749 *wwls24*[acdh-1p::GFP +unc-119(+)] MGH266 *rict-1*(*mg451*), VC222 *raga-1* (*ok386*), MGH559 *aak-2*(*ok754*);*zls356*[*daf-16p*::*daf-16a/b*::GFP+*rol-6*(*su1006*)], MGH249 *alxls19* [*bigr-1*::*bigr-1*::mRFP3-HA;*myo-2p*::GFP], WBM392 *Is(Pacs-2::GFP+rol-6(su1006))*, MGH430 *rsks-1(alx48* humanized S6K hydrophobic motif).

### *E. coli* Strains

Non-RNAi experiments were all conducted on NGM plates containing *E. coli* OP50-1 (CGC) as the food source and used 3-7 days after seeding. Cultures of *E. coli* OP50 were grown in Luria Broth (LB) for 15 hrs. at 37°C without shaking and seeded directly onto NGM plates. RNA interference experiments were conducted using *E. coli* HT115(DE3) bacteria (Ahringer library) as the food source. Clones were isolated from the primary RNAi library and plated on ampicillin/tetracycline plates. Individual clones were grown in LB broth for 15 hours with shaking. Cultures were concentrated 1:5 and seeded directly onto NGM plates containing. 5 mM isopropyl-B-D-thiogalactopyranoside and 200 mg/ml carbenicillin. Plates were used 1-5 days after seeding. All RNAi clones were sequence verified prior to utilization. Riboflavin solution or vehicle was applied to the plate and allowed to dry for at least 30 minutes prior to seeding with animals.

### Riboflavin Treatment

Culture grade riboflavin (Sigma-Aldrich) was dissolved in 50mM NaOH to a concentration of 13.3 mM (5 mg/ml). Fully seeded plates were treated with 500ul riboflavin solution (final concentration 665 μM) and allowed to dry on the plate prior to worm placement. Vehicle plates were treated with 500ul 50mM NaOH solution.

### Longevity assays

Lifespan analysis was conducted at 20°C except where indicated. Gravid adults were grown on NGM plates and isolated eggs were incubated overnight in M9 solution for hatching. Synchronized L1 animals were seeded unto RNAi plates and allowed to grow until the adult stage. Adult animals were subsequently transferred to fresh RNAi plates every other day until post-reproductive stage where they were maintained on a single plate. Dead worms were counted daily or every other day. Statistical analysis for survival curves was performed using OASIS2 software (Han, Lee et al. 2016).

### Development Assays

Synchronized L1s were prepared via bleach prep and plated on RNAi plates containing empty vector or *rft-1* RNAi grown at 20°C. Larvae were examined every 2 hours starting 41 hours after drop and scored for their transition to adulthood by the appearance of the vulvar slit.

### Brood Size

Fifty synchronized L1 animals of each strain were dropped on EV and *rft-1* RNAi treated plates. Two days later, 2 young adult animals from each condition were dropped onto new EV and rft-1 RNAi plates respectively. These animals were transferred every 2 days until the two adults from each condition became post-reproductive. All animals on residual plates were counted once they reached L4/YA to calculate brood size.

### Pharyngeal Pumping

Pumping rate was determined using a Sony camera attached to a Nikon SMZ1500 microscope that recorded 10 well fed Day 1 adult animals per genotype. Pharyngeal contractions in 15 second time periods for 4 technical replicates were counted (by slowing video playback speed by 4x) for each animal using OpenShot and pumping rates per minute were calculated.

### Oil-Red-O and Nile Red Staining

Lipid staining protocol was adapted from Escorcia et al (Escorcia, Ruter et al. 2018). Adult day 1 animals were collected via washing and washed twice with M9 solution. Animals were then fixed with 40% isopropanol for 3 minutes with shaking. For oil-red-O staining, working solution of oil-red-O was generated from stock solution and fixed animals were stained in working solution for two hours. Animals were subsequently placed in M9 solution to remove excess stain and were imaged on a Leica Thunder multichannel microscope to generate composite images. For Nile red staining, Nile red working solution was generated by mixing 6 ul/Nile red stock solution in 1 ml isopropanol. Animals were stained in working solution for two hours followed by 30 minutes of wash in M9 solution. Nile red imaging was performed on the Leica Thunder GFP setting with 10ms exposure at 5X magnification.

### Western Blotting

Worms were isolated by washing with M9 buffer and centrifuged into a pellet. Worm lysates were prepared by adding RIPA buffer and proteinase inhibitor cocktail (Roche) followed by water bath sonication in a Diagenode Bioruptor XL 4 at maximum strength for 15 minutes. Lysates were cleared of debris via centrifugation at 21,000g for 15 minutes at 4°C and supernatant was collected. Protein concentration as measured using the Pierce BCA Assay (Thermo Fisher). Lysate was subsequently mixed with 4X Laemmli buffer (Bio-Rad) and boiled for 10 minutes. Samples were run on SDS-PAGE protocol (Bio-Rad) and transferred to nitrocellulose membrane via wet transfer at 100V for 1 hour. Immunoblotting was performed according to primary antibody manufacturer’s protocols. Secondary antibody treatment utilized goat -anti-rabbit HRP conjugate or goat-anti-mouse-HRP conjugate (GE Healthcare) at 1:10,000 and 1:5000 dilutions, respectively. HRP chemiluminescence was detected via West-Pico substrate (Thermo Pierce). The western blot results shown are representative of at least two experiments. Primary antibodies used were the following: Rabbit monoclonal anti-Phospho-AMPKα (Thr172), Cell Signaling Technology Rabbit monoclonal anti-p70 Phospho-S6 Kinase (Thr389), Cell Signaling Technology Mouse monoclonal anti-Actin, Abcam

### Quantitative RT-PCR

Worms samples were flash frozen in liquid nitrogen and kept in -80°C until RNA preparation. Samples were lysed through the use of metal beads and the Tissuelyser (Qiagen) Total RNA was extracted using RNAzol RT (Molecular Research Center) according to manufacturer instructions. RNA was treated with RNAse free DNAse prior to reverse transcription with the Quantitect reverse transcription kit (Qiagen). Quantitative RT-PCR was conducted in triplicate using a Quantitect SYBR Green PCR reagent (Qiagen) following manufacturer instructions on a Bio-Rad CFX96 Real-Time PCR system (Bio-Rad) Expression levels of tested genes were presented as normalized fold changes to the mRNA abundance of control genes indicated in the figures by the δδCt method. The primers used for the qPCR are as follows:

*rft-1* forward: GCTATTGTTCAGATCGCGTGC

*rft-1* reverse: CAGAGACCCAATTGACAAATACATGC

*rft-2* forward: CGGGAGTTGTTCAGATCGCT

*rft-2* reverse: GAGTCCCAGTTGACAACAGCA

*rfk* forward: TGTTGGAAAAAGAAACGAAAGAA

*rfk* reverse: TCGATTAAAATTCGGTAACAACG

*flad-1* forward: TGCCTGGAGTTCCAAAATTC

*flad-1* reverse: GAAGGGCTGGGTGTTTTACA

*C07G1*.*7* forward: GCTGAAGAAGCTTCAACCGTAG

*C07G1*.*7* reverse: TCTCGTGTCAATTCCGGTCT

*hrg-9* forward: TGGAATATTGAGTGGCGTTG

*hrg-9* reverse: CCTCCTCTACTTGGTGCATGT

*cdr-2* forward: CGAGCCTCATTTGGAAAGAA

*cdr-2* reverse: GCATCTGCCGCTGTAACTTT

### GC/MS Lipid Analysis

Lipid extraction and GC/MS of extracted, acid-methanol derivatized lipids was performed as described previously (Pino and Soukas 2020). Briefly, 10,000 synchronous mid-L4 animals were sonicated with a probe sonicator on high intensity in a microfuge tube in 100-250 microliters total volume. Following sonication, lipids were extracted in 3:1 methanol: methylene chloride following the addition of acetyl chloride in sealed borosilicate glass tubes, which were then incubated in a 75°C water bath for 1 hour. Derivatized fatty acids and fatty alcohols were neutralized with 7% potassium carbonate, extracted with hexane, and washed with acetonitrile prior to evaporation under nitrogen. Lipids were resuspended in 200 microliters of hexane and analyzed on an Agilent GC/MS equipped with a Supelcowax-10 column as previously described (Pino, Webster et al. 2013). Fatty acids were indicated as the normalized peak area of the total of derivatized fatty acids detected in the sample, normalized by recovery of spiked-in, standard phospholipid and triglyceride.

### LC/MS Metabolite Analysis

Four biologic replicates of adult day 1 wild-type worms treated either with empty vector or *rft-1* RNAi were collected with M9 wash and frozen by liquid nitrogen into a worm pellet. Polar metabolites of homogenized worms were analyzed using a Thermo QExactive orbitrap mass spectrometer coupled to a Thermo Vanquish UPLC system, as previously described.(Garratt, Lagerborg et al. 2018) Bioactive lipids metabolites were profiled on the same system, as previous described.(Lagerborg, Watrous et al. 2019) Collected data were imported into the mzMine 2 software suite for analysis (version 2.53). Metabolites were annotated by using an in-house library of commercially available standards. Please see supplemental methods for detailed methods. All mass integration values, normalized abundance values, significance testing scores, and pathway enrichment scores are included in this manuscript as **Supplementary Table 2**.

### Quantification and statistical analysis

All western blotting quantifications were conducted on Bio-Rad Image Lab. Intensity analysis for fluorescent images was performed on ImageJ. Statistical analyses were performed using Prism (GraphPad Software). The statistical differences between control and experimental groups were determined by two-tailed students *t*-test (two groups), one-way ANOVA (more than two groups), two-way ANOVA (two independent experimental variables), or log-rank (survival analyses) as indicated in each figure legend, with numbers of samples indicated and corrected *P* values < 0.05 considered significant.

### Fluorescent Microscopy

DIC, brightfield and fluorescence Imaging of animals was performed utilizing the Leica Thunder microscopy system. Animals were picked onto a slide containing agar and 2.5mM levimasole solution. Imaging was performed within 5 minutes of slide placement. GFP and RFP Images were taken at 10ms exposure at 30% FIM and at 5X magnification, unless otherwise specified. Fluorescence intensity for quantification was calculated utilizing ImageJ software. For signal intensity experiments, quantification was performed on 20 worms (10 worms of two biological replicates).

### Two-Photon and Fluorescence Lifetime Imaging

Wild type and mutant *C. elegans* were immobilized for fluorescence imaging using a previously proposed protocol (Kim, Sun et al. 2013). Endogenous two-photon excited fluorescence (TPEF) images of *C. elegans* were acquired using a laser scanning microscope (Leica TCS SP8, Wetzlar, Germany) equipped with a femtosecond laser (Insight Deep See, Spectra Physics, Mountain View, CA). Fluorescence lifetime images (512 × 512 pixels) of C. elegans were acquired using the same excitation and emission settings as for intensity NAD(P)H and FAD images and a PicoHarp 300 time-correlated single photon counting unit (PicoQuant, Berlin, Germany) integrated in the Leica SP8 system. Please see Supplemental methods for further details on methods and analysis.

### Stimulated Raman Scattering imaging

Stimulated Raman scattering (SRS) images of C. elegans were acquired using a laser scanning confocal microscope (Leica TCS SP8, Wetzlar, Germany) equipped with a picosecond NIR laser (picoEmerald, APE, Berlin, Germany). SRS images were acquired in the wavenumber range of 2798 cm^-1^ to 3103 cm^-1^ with an interval of 6 cm^-1^. The Nd:VAN 1064.2 nm output was used as the SRS Stokes beam and the OPO beam tuned from 800 nm to 820 nm with step size of 0.4 nm was used as the pump laser. SRS images (620 × 620 microns x 51 wavenumbers, 512 × 512 pixels x 51 wavenumbers, 0.75 zoom) were acquired using a water immersion objective HCX IRAPO L 25x/0.95 NA with pixel dwell time of 4.9 μs. The pixels corresponding to regions occupied by C elegans were identified by implementing a global threshold of 300 (intensities ranged from 0 to 800). The SRS spectrum of each remaining pixel was normalized by the maximum SRS value of the entire field spectral image. To estimate the relative unsaturation level of the lipids in C. elegans, a ratio metric approach was adapted (Verma and Wallach 1977, Freudiger, Min et al. 2011, Nieva, Marro et al. 2012, Potcoava, Futia et al. 2014). Specifically, the ratio of the area under the SRS spectrum for wavenumbers spanning 2991 and 3022 cm-^1^ and wavenumbers spanning 2830 and 2870 cm-1, was estimated to represent the relative unsaturation levels (Verma and Wallach 1977, Freudiger, Min et al. 2011, Nieva, Marro et al. 2012, Potcoava, Futia et al. 2014). Both fluorescence and SRS images were calibrated for laser power before analysis.

## Acknowledgements

We thank Dr. Yuyao Zhang and Talia Hart for their creative input. This work was funded by the NIH/NIA Grants R01AG058259 and R01AG69677 and the Weissman Family MGH Research Scholar Award (to Alexander Soukas). This work was also funded by the Foundation for Women’s Wellness, the Human Growth Foundation and NIH/NIDDK F32DK124948 (to Armen Yerevanian). Thanks to the Nutrition Obesity Research Center (NORC) at Harvard (P30DK040561) for core services. Some strains were provided by the CGC, which is funded by the NIH Office of Research Infrastructure Programs (P40OD010440). Figure 3f and 6f were generated by BioRender.com. SRS and two photon images were acquired using microscopes acquired through NIH S10 OD021624 and NSF Major Research Instrumentation 1531683 grants.

## Author Contributions

Conceptualization: AY, LM, DB, AAS; Methodology: AY, LM, DB, SE; Validation: AY, LM, DB, SE; Investigation: AY, LM, SE, DB, SL, YZ, AA, LC, AAS; Formal Analyses: AY, EG, KD, MJ, IG, AAS; Writing: AY, AAS; Review and Editing: AY, FA, SE, LM, YZ, SL, LC, AA, DB, EG, MJ, IG, AAS; Funding: AY,IG AAS.

## Data Availability Statement

The data that support the findings of this study are available from the corresponding author upon reasonable request.

