## Supplemental Figures for "Riboflavin Depletion Promotes Longevity and Metabolic Hormesis in *Caenorhabditis elegans*"

**Supporting Information**

**Figure S1:** RT-PCR of riboflavin metabolism genes, DAPI staining of germline

**Figure S2:** DAF-16::GFP images, *sod-3p*::GFP images

**Figure S3:** FLIM Methods, Metabolite pathway analysis

**Figure S4:** Example images of *hsp-6p*::GFP activation

**Figure S5:** GC/MS lipid profiles of EV and *rft-1* RNAi treated animals

**Appendix S1:** Supplemental Methods

Supplemental References

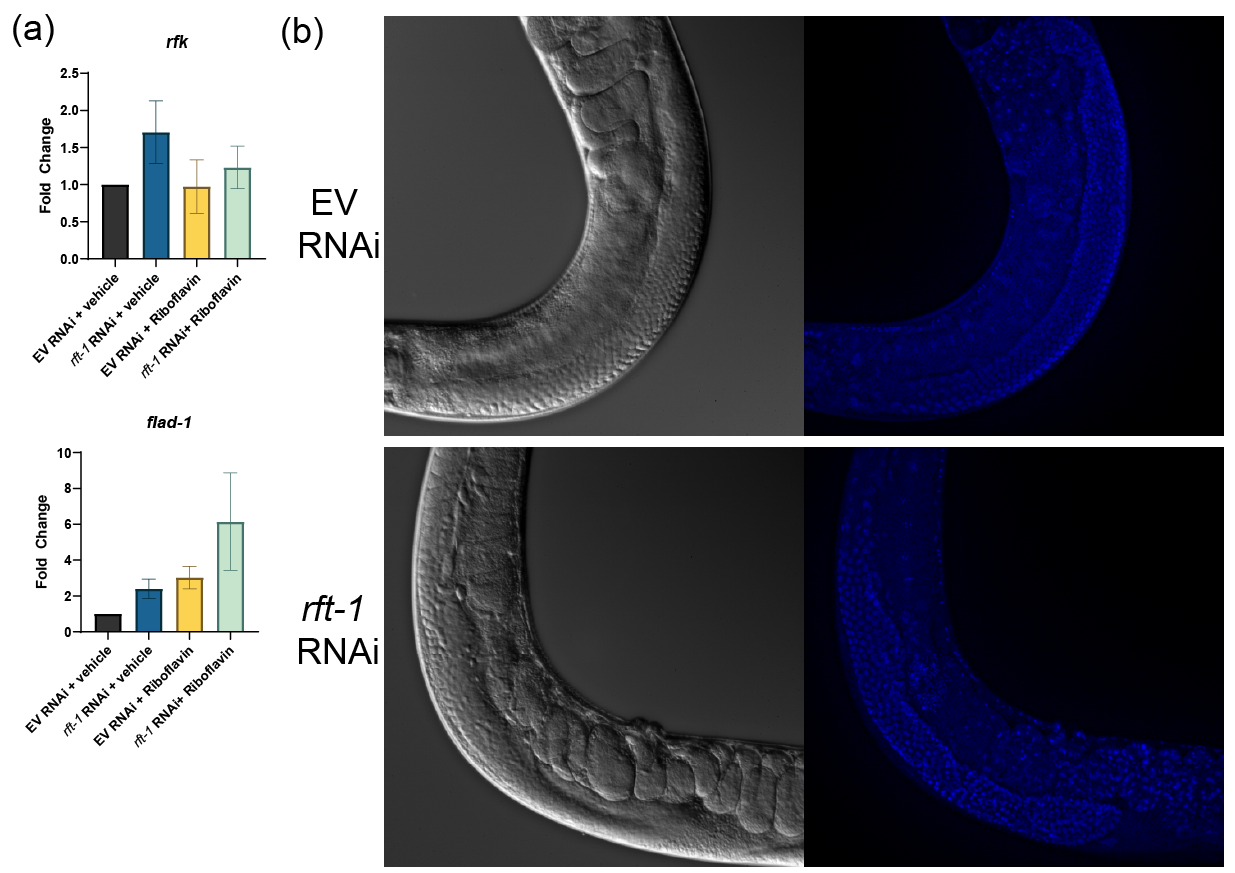

**FIGURE S1** (a) Quantitative RT-PCR of downstream flavin processing genes reveals no change in riboflavin kinase (*rfk*) and FAD synthetase (*flad-1*) levels with *rft-1* knockdown. Addition of riboflavin upregulates *flad-1* expression under both EV and *rft-1* knockdown. Results represent three biologic replicates. Bars represent means ± SEM. (b) Differential interference contrast (DIC) and fluorescence imaging of DAPI stained EV and *rft-1* RNAi treated adult day 1 animals reveals similar germ line stem cell morphology and the presence of oocytes.

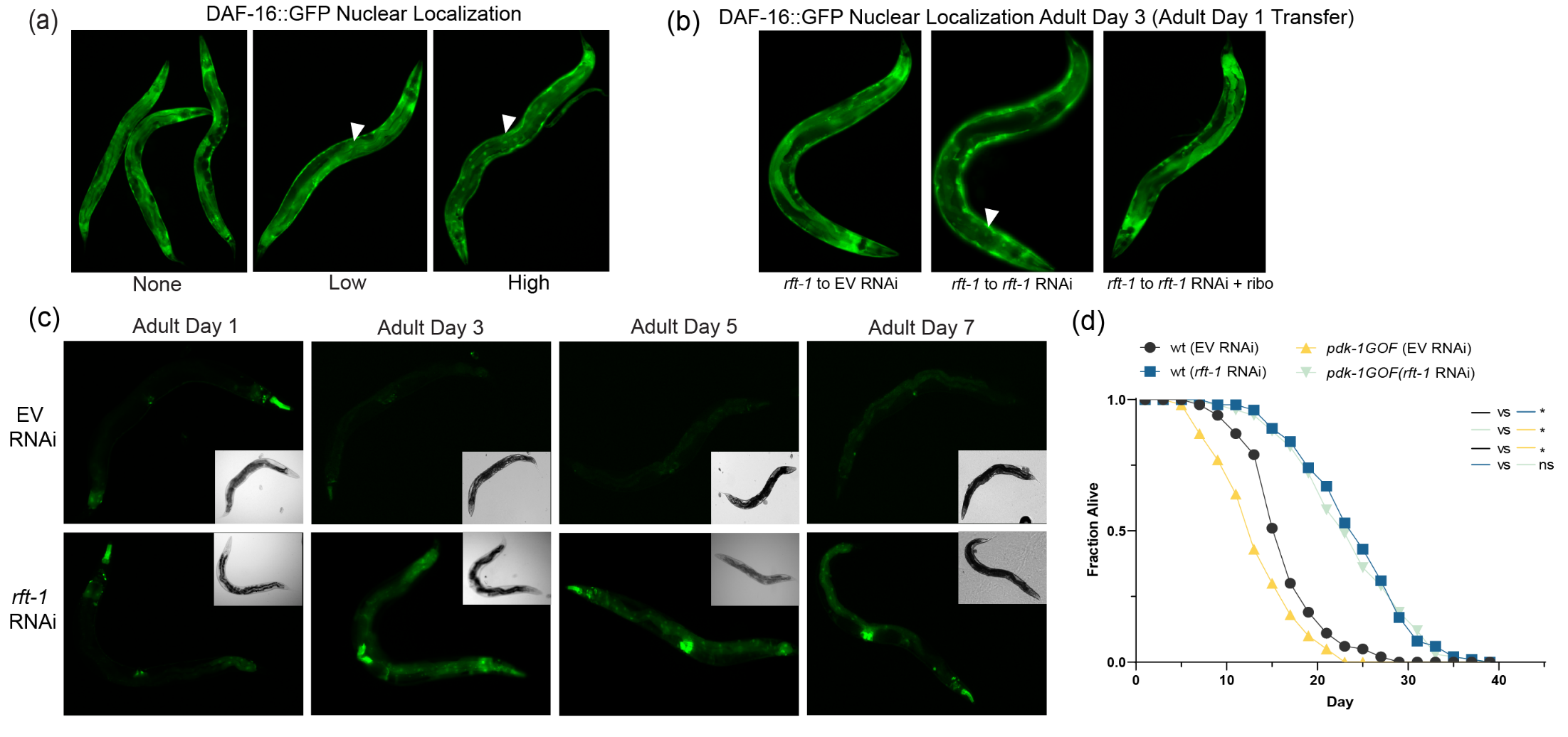

**FIGURE S2** Activation of DAF-16 by riboflavin deficiency. (a) Representative images of DAF-16::GFP high, low, and no (none) nuclear localization at 10X magnification (b) Transfer of *rft-1* RNAi treated animals at adult day 1 to empty vector (EV) RNAi plates or riboflavin supplemented plates shows that nuclear localization of DAF-16 is reversed by removal from *rft-1* RNAi and the addition of riboflavin. (c) Representative images of *sod-3p*::GFP animals over the lifespan, with significant increases evident from adult day 3 onward with *rft-1* RNAi. Persistent activation is seen in intestine, pharynx, and vulva. (d) Riboflavin depletion promotes lifespan extension in *pdk-1* gain of function animals. * indicates *P* < 0.05 by log rank analysis.

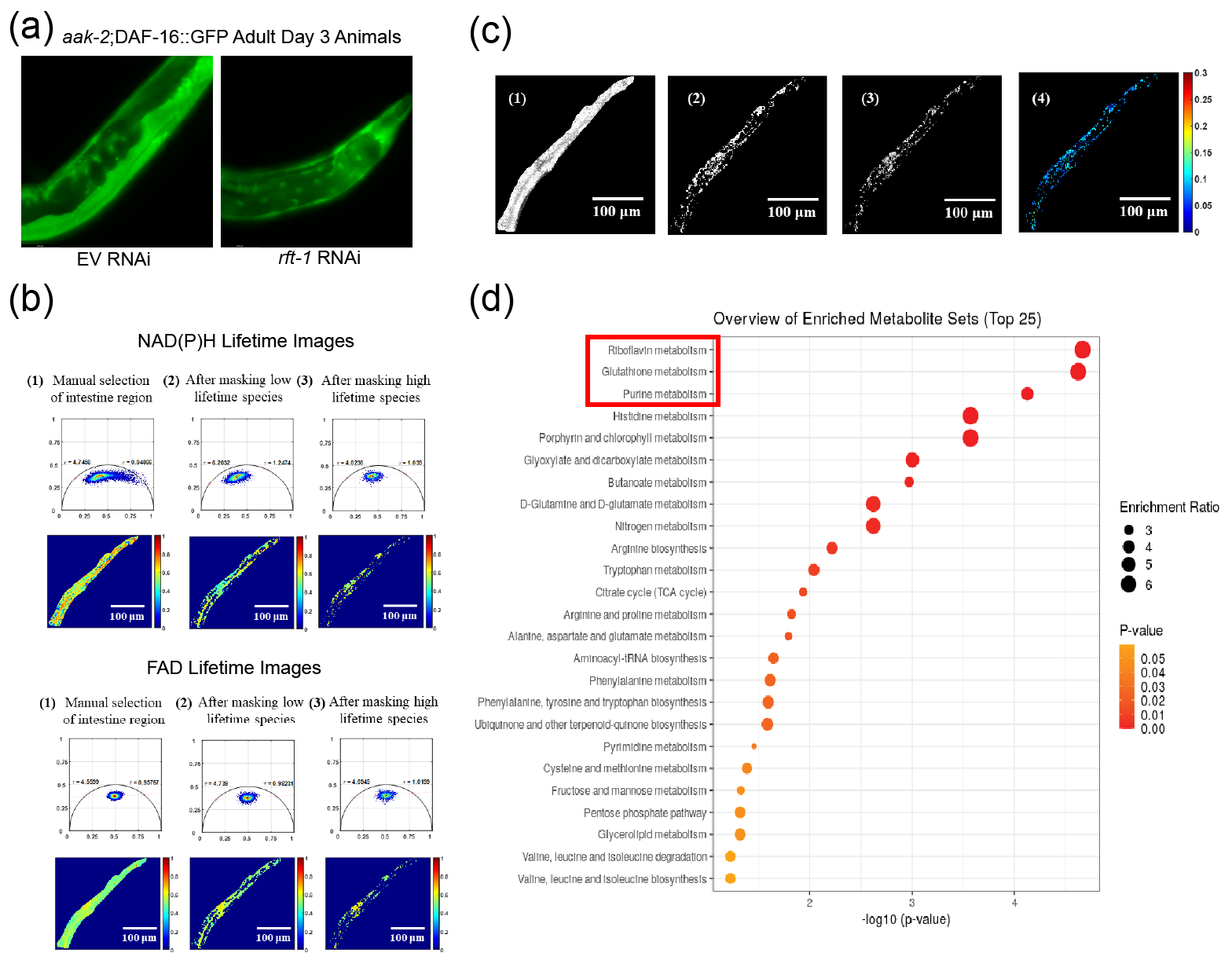

**FIGURE S3** Metabolite two-photon imaging and LC/MS quantification in riboflavin depletion. (a) Representative images of *aak-2*;DAF-16::GFP animals treated with EV and *rft-1* RNAi reveals nuclear localization with riboflavin depletion. (b) Representative phasors (*top panels*) and corresponding LLIF coded images (*bottom panels*) for NAD(P)H and FAD Lifetime Images. NAD(P)H images were acquired using 755 nm excitation/ 460 nm detection and FAD images were acquired using 860nm excitation/525 nm detection. (c) Representative images of total autofluorescence of NAD(P)H, FAD, and redox ratio map of *C. elegans* (1-4 respectively). Images of NAD(P)H, FAD, and redox map are shown only in masked region. (d) Summary plot of quantitative enrichment analysis (QEA) of metabolites between EV treated and *rft-1* RNAi treated animals reveals enrichment of pathways involved in riboflavin, glutathione and purine metabolism (n=4, P<0.01).

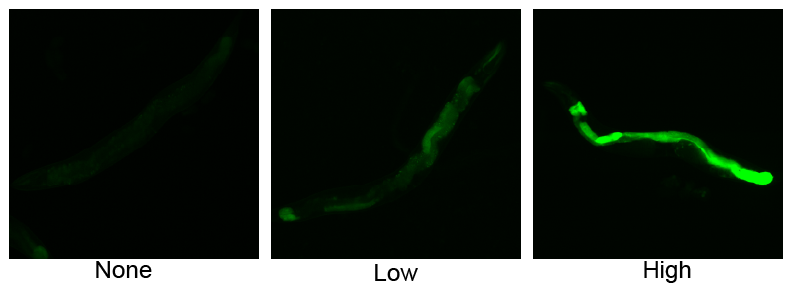

**FIGURE S4** Representative images of *hsp-6p*::GFP expression categories in Figure 4a.

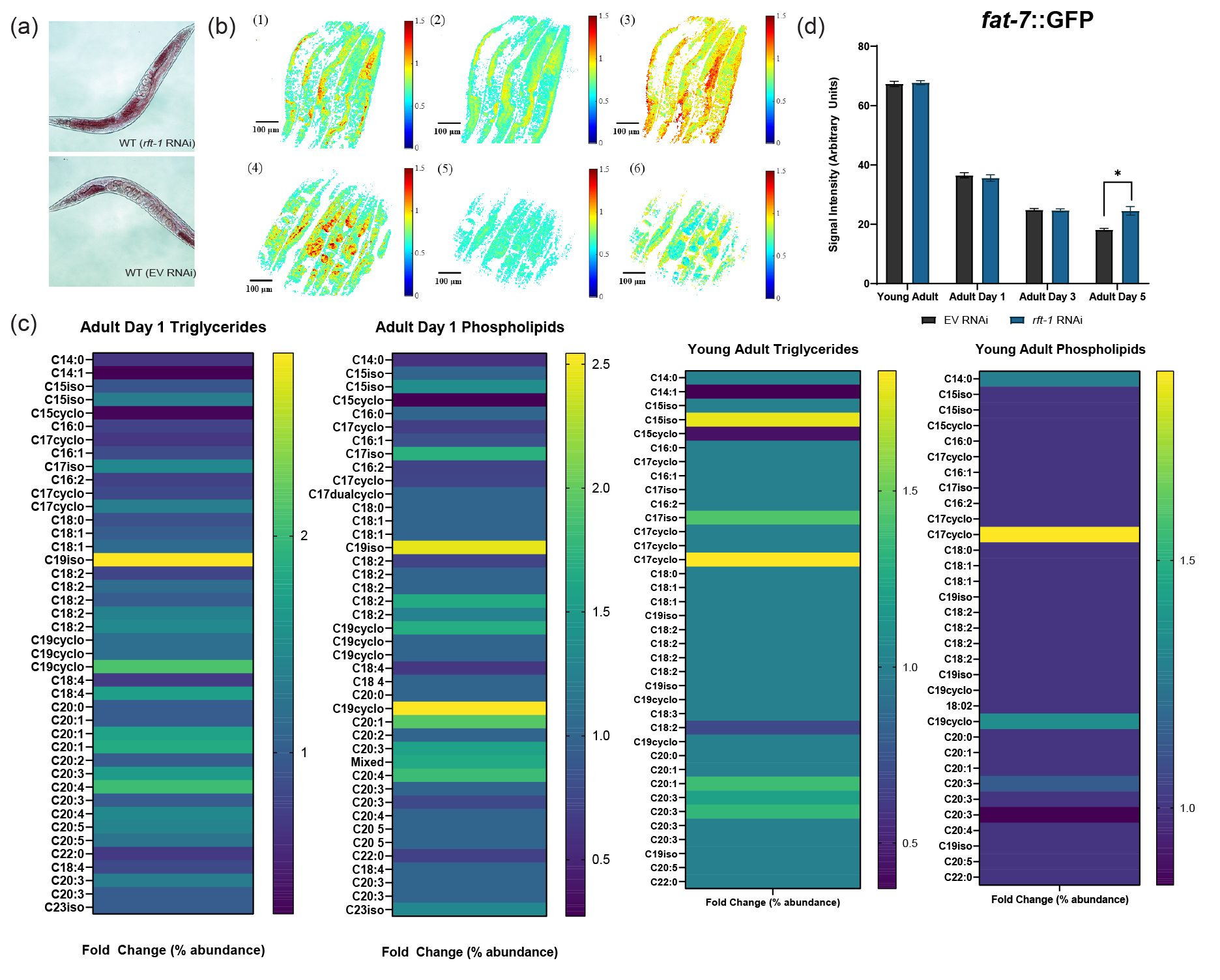

**FIGURE S5** Lipid analyses with riboflavin depletion indicate increased fat mass. (a) Oil-red-O staining of adult day 1 animals treated with EV and *rft-1.* (b) Pseudo color map of (1, 4) total unsaturated fatty acid (area under curve between wavenumbers 2991 and 3022 cm^-1^), (2, 5) total fatty acid (area under curve between wavenumbers 2830 and 2870 cm^-1^) and (3, 6) ratio of TUFA and TFA for EV treated (4-6) and *rft-1* RNAi treated (1-3) *C. elegans* acquired using SRS imaging. (c) GC/MS analysis of triglyceride and phospholipid fractions in adult day 1 and young adult worms. Ratios represent fold change of worms treated with *rft-1* RNAi compared to EV treated animals (d) Imaging of *fat-7p*::GFP animals reveals unchanged expression of *fat-*7 until adult day 5, where *rft-1* RNAi preserves an aging-related decrease in desaturase expression. * indicates P < 0.05 by two-tailed Student’s t-test (b) and by two-way ANOVA (c). Bars represent means ± SE.

**Supplemental Methods**

**DAPI Staining**

Adult day 1 animals were collected via washing and washed twice with M9 solution. Animals were then fixed with 40% isopropanol for 3 minutes with shaking. For DAPI staining, a working solution was generated by mixing 2ul of 1 mg/ml DAPI solution in 1ml of 40% isopropanol. Animals were stained in working solution for 45 minutes. Imaging was performed on the Leica Thunder A4 setting with 50ms exposure at 20X magnification.

**LC/MS Metabolite Analysis**

Each worm pellet contained 4000 worms. Worm pellets were transferred to 2 mL impact resistant homogenization tubes containing 300 mg of 1 mm zirconium beads and 1 mL of 80:20 ethanol:water containing internal standards(Lagerborg, Watrous et al. 2019). Using an Omni Bead Ruptor Elite, samples were homogenized in three 10 second cycles at speed 8 m/s with 10 seconds pause between cycles to prevent overheating. All samples were then transferred to a clean 1.5 mL microcentrifuge tubes and placed at -20 °C for 20 minutes to facilitate protein precipitation. Samples were then centrifuged at 14,000 g for 10 minutes at 4 °C. After which 150 uL of supernatant was processed through solid-phase extraction to isolate lipid metabolites using a Phenomenex Stata-X polymeric 10 mg/mL 96-well SPE plate, as previously described. For polar metabolites, 50 uL of supernatant was dried *in vacuo* and resuspended in 50 uL of 40:40:20 acetonitrile:methanol:water containing 1 ng/uL of L-Phenylalanine-13C9,15N.

All visualization and significance testing of metabolomics was conducted using the MetaboAnalyst 5.0 package (Pang, Chong et al. 2021). Mass integration values for 121 identified metabolites were extracted from full-scan LC-MS/MS measurements of L4 wildtype (Bristol N2) *C. elegans* treated with L4440 vector (EV) or *rft-1* RNAi throughout larval development (n = 4 independent biological replicates). Abundance values were subsequently log_10_ transformed and normalized by mean centering and division by the standard deviation of each variable (auto-scaling). Normalized abundance values were then assessed for statistical significance via independent two-sample t-tests followed by false discovery rate (FDR) control using the Benjamini-Hochberg (BH) method. Volcano plot visualizations were plotted using the -log_10_(FDR) in comparison to the log_2_(fold change) between normalized *rft-1* RNAi/EV using Prism 9 (GraphPad Software). Metabolites were considered differentially abundant with an FDR controlled *P* value < 0.05. The top 30 metabolites across treatment (ranked by *t* statistic and FDR value) were visualized using a heatmap, with hierarchical clustering of samples and normalized compound abundances included. Quantitative enrichment analysis (QEA) was conducted on differentially abundant metabolites using internal R package *globaltest*, calculating an average *Q* statistic across all compounds belonging to a given metabolic pathway as defined by the KEGG database (Goeman, van de Geer et al. 2004, Kanehisa, Sato et al. 2016). Dotplots were generated by comparing the -log_10_(FDR) with the *Q* enrichment ratio of the top 25 most enriched metabolite sets.

**Two-Photon and Lifetime Imaging**

C. elegans were mounted over 5–10% agarose pads on glass slides and 0.25–2 µL of a suspension of polystyrene beads (Polysciences, 2.5% by volume, 0.1 µm diameter) were applied around the sample and on the coverslip. The coverslip was placed upside down and worms were immobilized in gel-microbead matrix between glass slide and coverslip. The cover glass was sealed with nail polish to prevent dehydration.

As per our previous studies (Quinn, Sridharan et al. 2013, Varone, Xylas et al. 2014, Liu, Pouli et al. 2018), NAD(P)H images were acquired with an excitation wavelength of 755 nm and recorded at 460 ± 25 nm using a non-descanned detector. Fluorescence spectra of NADH and NADPH are indistinguishable, and their combined fluorescence is represented as NAD(P)H. FAD images were acquired with an excitation wavelength of 860 nm wavelength and recorded at 525 ± 25 nm using a non-descanned detector. Laser light was focused on the sample using a water immersion 40x objective with 1.1 numerical aperture and average laser powers of 24 mW for both wavelengths. The TPEF images (290 × 290 μm, 1024 × 1024 pixels, 1 zoom) were acquired with pixel dwell time of 0.7 μs and 8-line average.

For analysis of the intensity and lifetime TPEF images, the intestine region of the C. elegans was selected manually relying on the integrated intensity of the NAD(P)H lifetime images, since the autofluorescence signal was significantly higher than in the surrounding tissues. Pixel-wise lifetime data was analyzed using the phasor approach, as described previously (Liu, Pouli et al. 2018). Briefly, after Fourier decomposition at the repetition frequency of the laser, the normalized cos and sin components were represented along the x and y axis, respectively, so that the fluorescence lifetime spectrum from each pixel was represented by a single point in phasor space. The phasors of spectra corresponding to a single exponential decay were represented by points that lay on the universal semi-circle, while biexponential decays like that of NAD(P)H and FAD, were represented by ellipsoid shapes within the semicircle. A line fit to the major axis of such an ellipsoid crossed the universal semi-circle at points that corresponded to the short and long lifetime components. The relative distance of the ellipsoid centroid on the line represented the long lifetime intensity fraction (LLIF) of the fluorophore, which is correlated with the mean lifetime (Alonzo, Karaliota et al. 2016, Liu, Meng et al. 2021). The original phasors of *C elegans* presented more complex shapes indicating the presence of additional fluorophores. The corresponding fluorescence lifetime images coded according to the LLIF value revealed the presence of numerous granule-like structures with blue hues corresponding to very low LLIF values. Strong fluorescence from such granules has been previously reported and attributed to tryptophan metabolite anthranilic acid glucosyl esters or advanced glycation end products (Coburn and Gems 2013, Teuscher and Ewald 2018, Komura, Yamanaka et al. 2021). To remove contributions from these granules, all pixels yielding NAD(P)H LLIF values lower than 0.4 were identified and removed. Lipid droplets have also been identified in *C. elegans* intestinal cells. Since lipids can autofluoresce (Datta, Alfonso-García et al. 2015, Alonzo, Karaliota et al. 2016), we implemented an additional thresholding procedure established in our previous study to identify and remove such contributions (Alonzo, Karaliota et al. 2016). This method relied on the observation that lipid droplets within white and brown adipose tissues were characterized by high LLIF in the 755/460nm channel (Alonzo, Karaliota et al. 2016). Three level Otsu thresholding was applied to the images in the 755/460nm and 860/525 nm channel images to remove weakly autofluorescent nuclei and intercellular regions. Then the regions with LLIF of more than 0.7 were identified as lipid droplets and removed from the image. When contributions from the pixels removed based on low and high LLIF thresholding were also removed from the phasors associated with the fluorescence associated with 860 nm excitation/525 nm detection, there was limited impact on the shape of the phasor, indicating that lipid droplet fluorescence was not prominent in that wavelength regime, consistent with our previous findings (Alonzo, Karaliota et al. 2016). The regions that contained pixels with LLIF values below the low threshold and above the high LLIF threshold were dilated by applying a 5 × 5 pixel averaging window on the corresponding binary masks to ensure that borders from these organelles were excluded from the cytoplasmic region used to assess NAD(P)H and FAD contributions. The optical redox ratio, was calculated as the pixel-wise ratio of FAD/(NAD(P)H+FAD) intensity images. The intensity fluctuations with the NAD(P)H images were analyzed used approaches described in detail previously to assess the levels of mitochondrial clustering/fragmentation (Pouli, Balu et al. 2016, Liu, Pouli et al. 2018)
